# EDTA v2: enabling scalable TE annotation in animal genomes

**DOI:** 10.64898/2026.07.01.735963

**Authors:** Shujun Ou, Tianyu Lu, Hieu Nguyen, Kenji Gerhardt, Ning ‘Faye’ Fang, Usman Rashid, Joseph Guhlin, Jacques Dainat, Zhigui Bao, Philipp E. Bayer, Yeojung Na, Christopher Benson

## Abstract

The Extensive de-novo TE Annotator (EDTA) automates transposable element annotation in plant genomes but lacks direct LINE/SINE detection, limiting its applicability to animal genomes. We present EDTA v2, which integrates LINE and SINE detection, completely rewrites TIR-Learner for deployability and scalability, and accelerates structural detectors by up to two orders of magnitude. Tested in 30 animal genomes from the Vertebrate Genomes Project Phase I, EDTA v2 bridges the non-LTR detection gap that has prevented automated TE annotation in animals.

## Main text

Transposable elements (TEs) are a major component of eukaryotic genomes and play important roles in genome evolution, gene regulation, and speciation^1,2^. We developed the Extensive de-novo TE Annotator (EDTA) to automate whole-genome TE annotation by integrating structural and homology-based approaches into a single pipeline^3^. EDTA has since been widely adopted in plant genomics (e.g., refs^4–7^), with its detection scope primarily focused on LTR retrotransposons, TIR transposons, and Helitrons. Non-LTR retrotransposons (LINEs and SINEs), which dominate most animal genomes, were not specifically detected. Gozashti and Hoekstra^8^ highlighted this gap, noting that EDTA’s plant-centric design restricted its applicability to the growing number of animal genome assemblies. In particular, coordinated efforts as part of the Earth BioGenome Project are aimed at generating high-quality reference genomes for all named eukaryotic species in the next ten years. For vertebrates, the Vertebrate Genomes Project^9^ recently released its Phase I data freeze^10^. In addition to its restricted detection scope, the core structural detectors in EDTA v1 scaled poorly on large genomes. These genomes are also becoming increasingly available thanks to improved sequencing technologies and assembly algorithms^11^.

To address these limitations, we previously outlined a plan to expand EDTA for broader use in animal genomes^12^. Here we present EDTA v2, which introduces *de novo* LINE and SINE detection, a complete rewrite of TIR-Learner, and accelerated structural detectors (**Fig. 1A**). We integrated RepeatModeler2 (ref^13^) and AnnoSINE_v2 (ref^14^) in the initial detection stages for LINEs and SINEs (non-LTRs), respectively, with TEsorter^15^ providing accurate classification. The TIR detection module, TIR-Learner, was completely rewritten to remove brittle dependencies, eliminate heavy temporary I/O, add TIRvish^16^ as a second discovery engine alongside GRF^17^, and enable distribution as a single Bioconda recipe (**Fig. S1**). This resolved a longstanding deployment barrier where users routinely failed to install TIR-Learner on varying compute platforms or conda environments, making TIR annotation inaccessible for many groups. The TIR-Learner rewrite enables the annotation of highly fragmented assemblies for roughly 2× speedup without sacrificing annotation integrity (**Fig. 1B**, **Fig. S2, S3, S4**). We also wrapped LTR_HARVEST and HelitronScanner in parallelized accessory scripts, achieving 5 to 240× speedup on a modern server (**Fig. 1C,D**) and enabling annotation of genomes as large as the 29-Gb axolotl in less than two hours. To accommodate the expanded detection scope, a cross-category filtering system was developed to resolve misclassifications across all five TE subclasses through richness-ratio purification, hierarchical masking, and all-versus-all redundancy removal.

**Figure 1.**
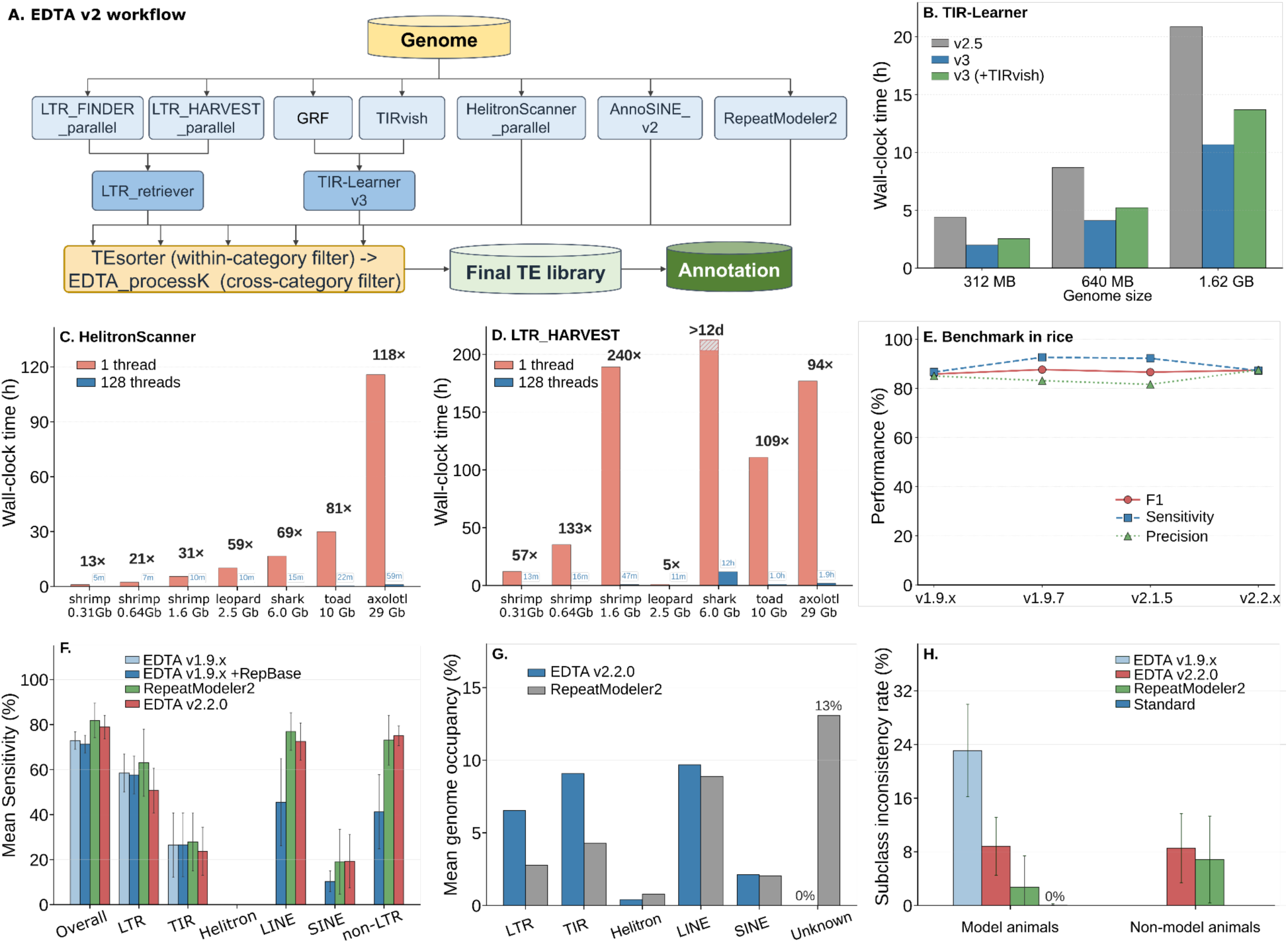
The EDTA v2 pipeline and benchmark performance on animal genomes. (**A**) EDTA v2 workflow. The input genome is processed in parallel by structure-based or *de novo* detectors for each major TE subclass. Raw libraries are filtered within and across subclasses, producing the final non-redundant TE library used for whole-genome annotation. Scalability of TIR-Learner v3 (28 threads) (**B**), HelitronScanner_parallel (**C**), and LTR_HARVEST_parallel (**D**), showing up to 240-fold speedup of the parallelized implementations. Subsamples of the highly fragmented Pacific white shrimp genome (contig N50 87.4 kb)^21^ were used. (**E**) Performance of EDTA across major releases on the rice genome. (**F**) Mean sensitivity on five animal models (chicken, fly, human, mouse, and zebra finch) for which curated TE libraries are available. All EDTA versions were run in *de novo*-only mode, except EDTA v1.9.x +RepBase, which was supplemented with curated TE libraries from RepBase. (**G**) Per-category mean genome coverage across 30 non-model animal genomes for EDTA v2 (blue) and RepeatModeler2 (grey). Tandem repeats were removed from the Unknown category. (**H**) Mean classification-inconsistency rate across TE subclasses on five model genomes (chicken, fly, human, mouse, zebra finch) and 30 non-model animal genomes. Standard refers to the curated reference library of each model species. Error bars are standard deviation across species.

**Figure 2.**
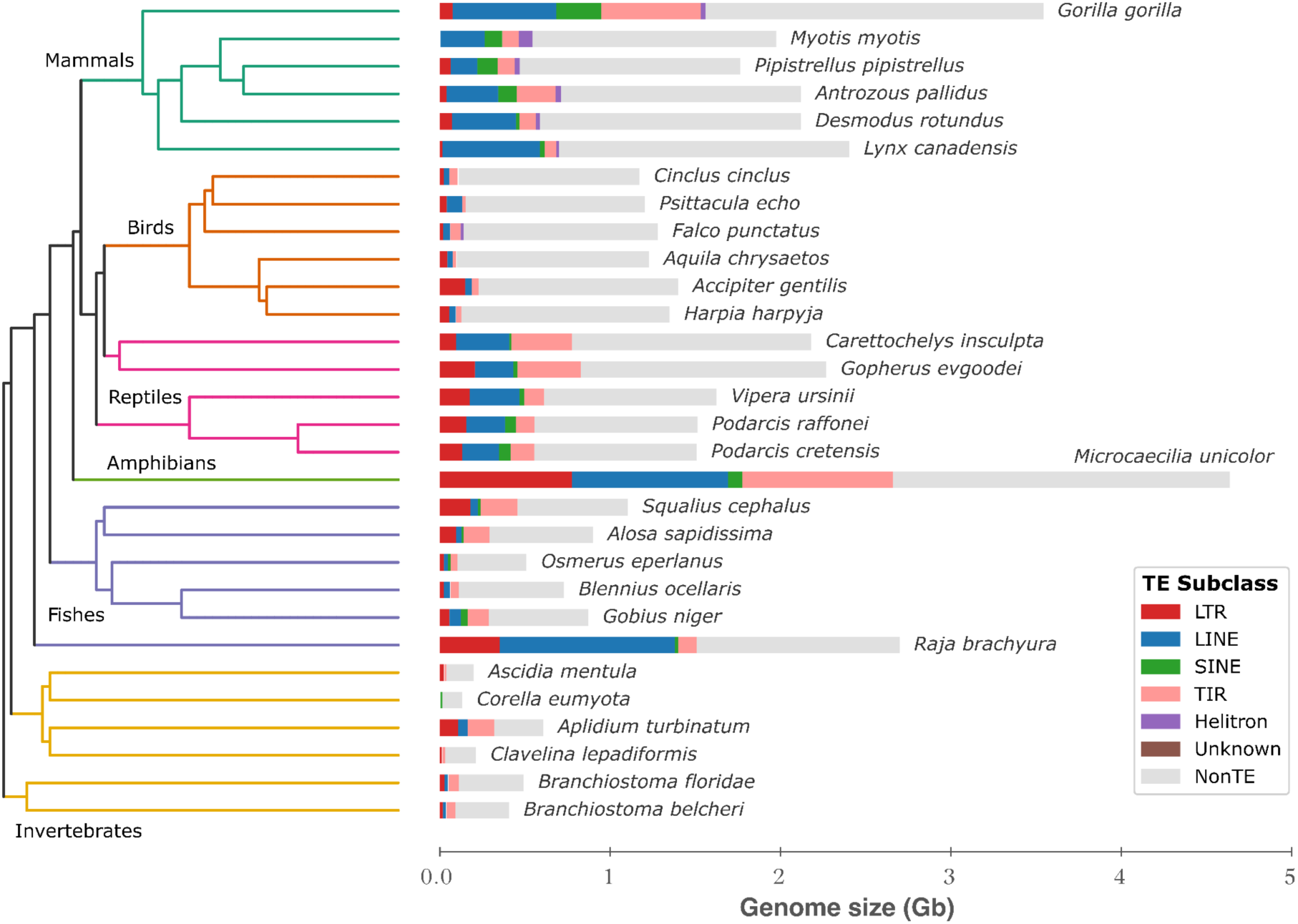
Transposable element composition across 30 animal species. The rectangular cladogram has branches colored by major clade, with clade names annotated. Stacked horizontal bars show the proportion of each genome annotated as TEs at the subclass level. Bar width is proportional to assembly size.

On the rice and maize genomes, EDTA v2 matches v1 performance. Total-annotation sensitivity, precision, and F1 are all maintained at 0.87-0.93 while raising non-LTR F1 from zero to 0.63-0.68 (**Fig. 1E, Fig. S5, S6**). These results demonstrate that the expanded detection scope adds non-LTR sensitivity without compromising existing TE categories in plants.

On the five animal genomes for which curated TE annotations are available (chicken, fly, human, mouse, and zebra finch), EDTA v2 attained total sensitivity and F1 scores of 0.79 (SD = 0.12) and 0.75 (SD = 0.05), respectively (**Fig. 1F, Fig. S7**). The largest gains were in non-LTR retrotransposons, where mean non-LTR F1 rose from zero (EDTA v1) to 0.76 (**Fig. S7**). Further, the EDTA v2 *de novo* annotation outperforms EDTA v1 with supplemental RepBase libraries in both sensitivity and F1 score (**Fig. 1F, Fig. S7**), demonstrating that the integrated structural detection surpasses database-dependent annotation. EDTA v2 also attains the highest F1 in SINE detection among all tested tools, reflecting the benefit of dedicated AnnoSINE_v2 integration. In these model genomes, RepeatModeler2 achieves higher overall F1 scores by using curated TE databases for classification and validation (**Fig. 1F, Fig. S7**).

We benchmarked EDTA v2 on 30 animal genomes from the Vertebrate Genomes Project Phase I, including 24 vertebrates and 6 invertebrate outgroups. EDTA v2 left a near-zero fraction of unclassified TEs, whereas RepeatModeler2 produced an average of 13% Unknown repeat content per assembly (**Fig. 1G**) and up to 36.7% in *Microcaecilia unicolor* (**Fig. S8, S9**). Furthermore, unclassified repeats averaged 47.2% (SD = 18.0%, tandem repeat purged) of all RepeatModeler2-annotated repeats in non-mammalian species (**Fig. S8; Table S1**), which is likely due to their lack of sequence similarity to the mammalian-centric database. This contrast reflects EDTA v2’s structural-classification design, where TE candidates are identified by structural features shared between divergent species (**Fig. 1A**).

EDTA v2 produces comparable or higher occupancy estimates for LTR, LINE, SINE, and TIR elements and a near-zero Helitron fraction in the tested animals (**Fig. 1G**), consistent with the known scarcity of autonomous Helitrons outside plants and a few bat lineages^18^. Finally, total annotated TE fractions are comparable between the two tools (**Fig. S8**), with 31.8% of Unknown TEs reported by RepeatModeler2 largely reclassified by EDTA v2 into TIR (37.6%) and LTR (20.7%) elements (**Fig. S9, S10**). We further evaluated the subclass-level annotation inconsistency that does not rely on curated annotations. Among animal model species, the inconsistency rate fell from 23.1% for EDTA v1 to 8.8% for EDTA v2 (**Fig. 1H**). On the 30 non-model genomes, annotation inconsistencies between EDTA v2 and RepeatModeler2 were statistically indistinguishable (paired t-test, *P* = 0.25). EDTA v2 thus held a stable inconsistency rate near 9% in both model and non-model animal species (**Fig. 1H**).

Consistent with established phylogenetic expectations, EDTA v2 recovered LINE-dominated landscapes in mammals (primarily L1) and sparse TE profiles in birds, while fish and reptile genomes showed more balanced contributions from DNA transposons and LTR retrotransposons (**Fig. 2**). Total TE content broadly correlated with assembly size, underscoring how lineage-specific amplification histories shape genome architecture. These patterns, recapitulated here entirely from *de novo* structural annotation, demonstrate that EDTA v2 produces biologically coherent TE landscapes previously inaccessible in automated *de novo* annotation (**Fig. 2; Fig. S10A**).

## Discussion

EDTA v2 closes the central gap that limited the original pipeline to plant genomes by integrating direct LINE and SINE detection and accelerating the rate-limiting structural detectors, making automated TE annotation tractable on multi-gigabase animal assemblies. Across 30 animal genomes, the pipeline produces nearly complete superfamily-level classifications in contrast to RepeatModeler2, which, on average, presents 47.2% of annotated repeats as Unknown in non-mammal species. On model genomes for which curated reference libraries exist, RepeatModeler2 still achieves higher F1 because its *de novo* families can be matched to well-annotated databases. For the much larger universe of non-model genomes where curated libraries of TEs are absent, EDTA v2’s structurally anchored classification provides immediate, interpretable annotations that can serve as a starting point for comparative genomics and downstream curation (e.g., Dfam and Repbase^19^). An accuracy gap remains compared to expert curation. Towards fully curated libraries, a natural next step for EDTA v2 is extension and edge refinement by including tools such as TEtrimmer^20^, which automates previously manual curations. EDTA v2 makes automated, classified TE annotation accessible across the growing catalogue of non-model animal genomes.

## Software availability

EDTA v2.2.0 is freely available at https://github.com/oushujun/EDTA under the GPL-3.0 license. The program is installable via Bioconda and is distributed as OCI container images (Docker and Singularity). TIR-Learner v3 is available at https://github.com/lutianyu2001/TIR-Learner, and is likewise released under the GPL-3.0 license, available through Bioconda, and distributed as OCI container images (Docker and Singularity).

## Author Contributions

S.O. conceived the study, designed, and implemented EDTA v2 and LTR_HARVEST_parallel. T.L. designed and implemented TIR-Learner v3. K.G. designed and implemented HelitronScanner_parallel. H.N. annotated the animal genomes. S.O., T.L., K.G., N.F., U.R., J.G., J.D., Z.B., P.B., Y.N., and C.B. contributed to coding, testing, debugging, and making conda recipes. S.O., T.L., and K.G. performed benchmarks of various tools. S.O. wrote the manuscript, and all authors commented and approved the final version.

## Acknowledgements

We thank Antonio Gu, Qiushi Li, Sergei Ryazansky, Nick Carleson, Sanzhen Liu, Zhougeng Xu, Shun Wang, Nancy Manchanda, Eric Burgueño, and many others for testing, debugging, and improving the EDTA pipeline. S.O. was supported in part by the OSU STEM Education and JobsOhio award. C.B. was supported by the USDA National Institute of Food and Agriculture AFRI Postdoctoral Fellowship (#2024-67012-43408). EDTA benchmarking and genome annotation were supported in part by NSF ACCESS Discover grants BIO250178 and BIO250194 to S.O. We thank the Purdue Rosen Center for Advanced Computing for providing technical support to the Anvil cluster.

## Competing Interests

The authors declare no competing interests.

## Supplementary Figures

**Figure S1.**
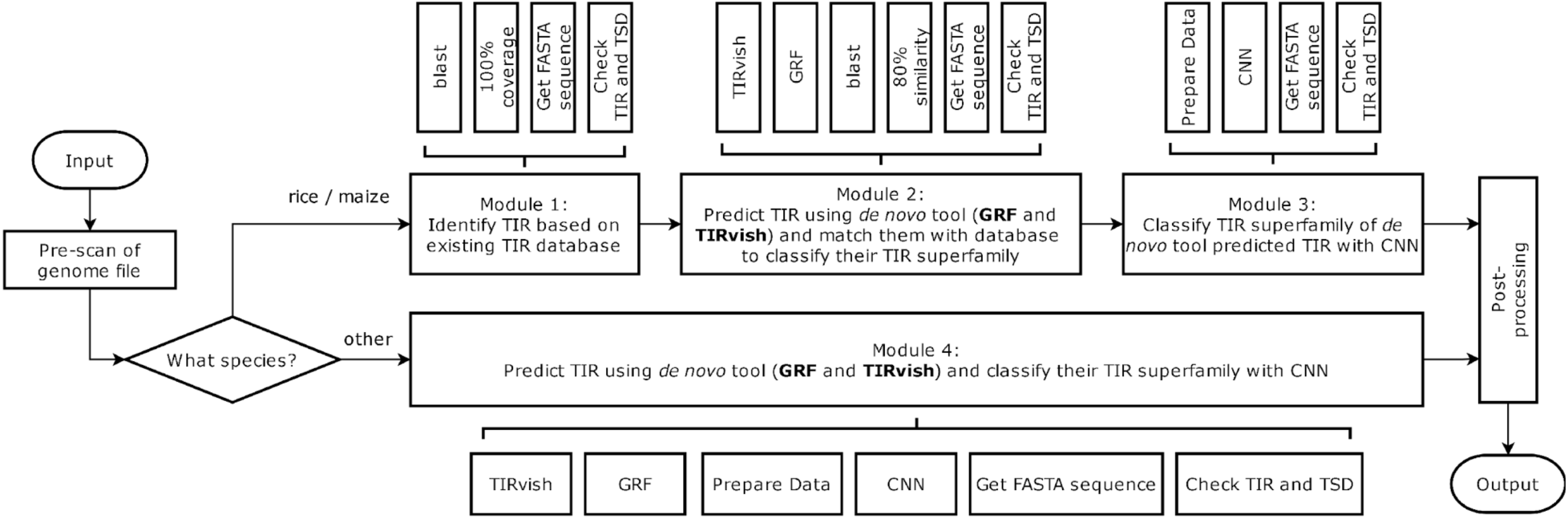
Workflow of TIR-Learner v3.

**Figure S2.**
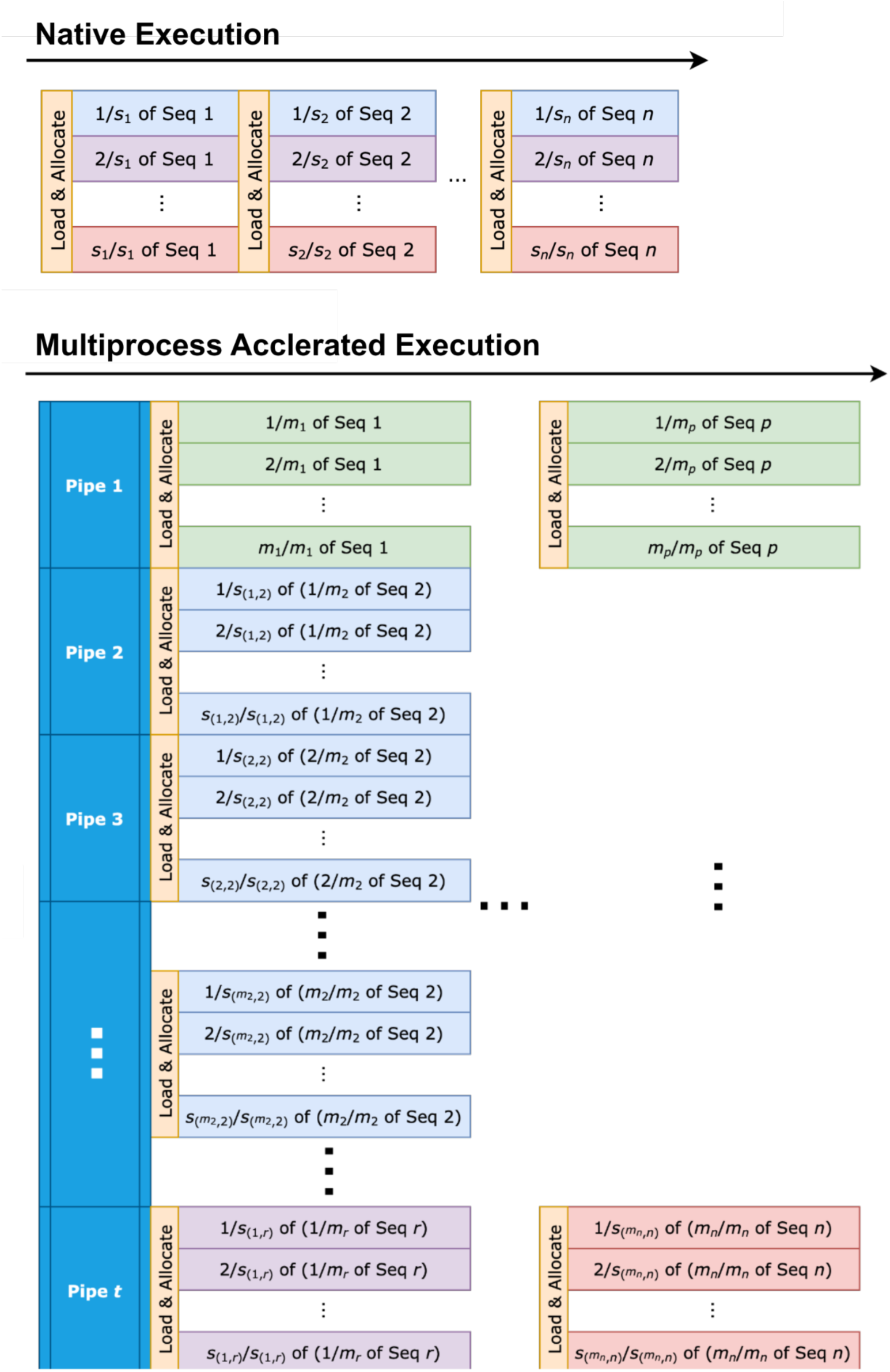
Accelerating GRF with improved load balancing. Compared with GRF’s native multithreading, our hybrid parallelization approach wraps multithreading in an outer multiprocessing layer in which the available CPUs (*C*) are partitioned into *P* = ⌊√*C*⌋ concurrent processes, each allocated *T* = ⌊*C*/*P*⌋threads. This balances two competing costs under a fixed core budget (*P* × *T* ≤ *C*): with many short input sequences, per-sequence loading overhead can exceed the actual analysis time, so concentrating all cores into native MP threads (*P* = 1) leaves serial loading as the dominant bottleneck, whereas excessive process counts incur scheduling overhead. Partitioning at *P* ≈ *T* ≈ √*C* minimizes *P* + *T*, parallelizing loading across processes while retaining sufficient threads per process for analysis.

**Figure S3.**
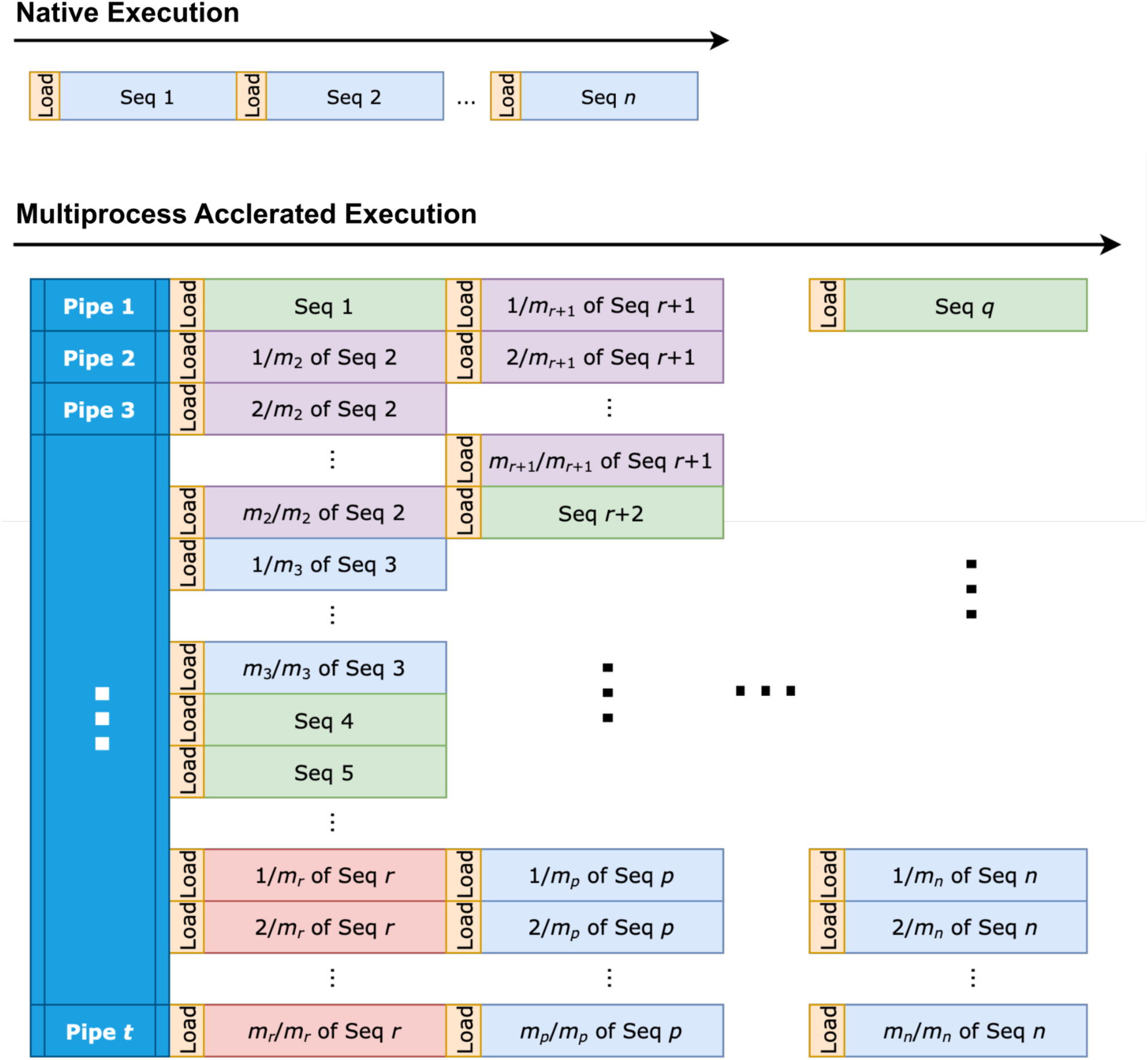
Accelerating TIRvish with improved load balancing.

**Figure S4.**
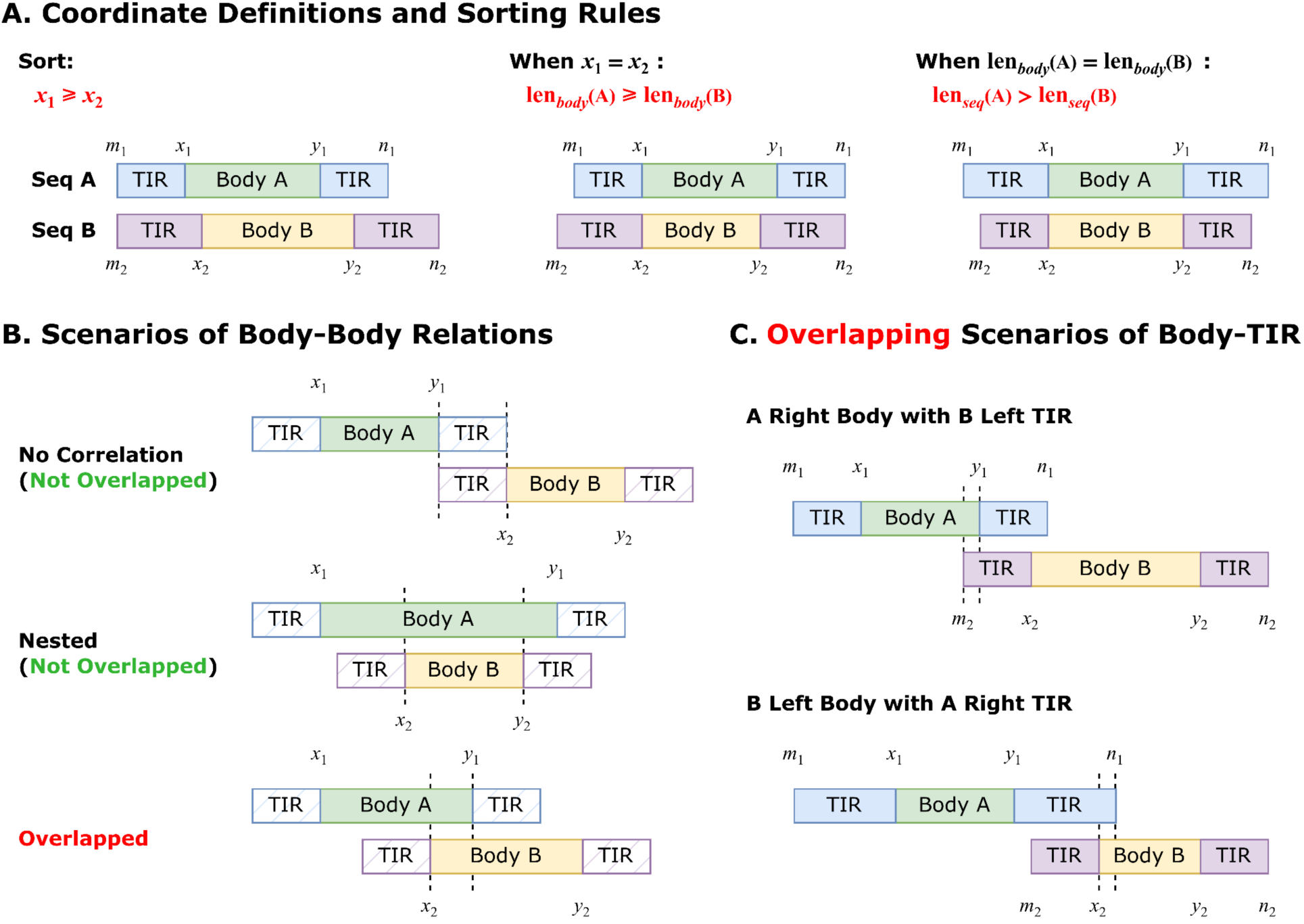
Deduplication criteria for candidates identified from overlapping genomic windows. (**A**) Coordinate definitions and sorting rules for two candidate TIR elements. For each element, *m* and *n* denote the outer boundaries (full-length element including TIRs), while *x* and *y* denote the inner boundaries of the main body (excluding TIRs). Candidates are sorted such that *x*_!_ ≥ *x*” ties are resolved by decreasing main body length, then by decreasing entire element length. (**B**) Body–body overlap scenarios based on main-body coordinates. Partial overlap between the main bodies of two elements is classified as overlapped (duplicate); complete nesting of one element body within another is classified as not overlapped (distinct). (**C**) Body–TIR overlap scenarios. When the main body of one element overlaps with the TIR of another element, the pair is classified as overlapped. Elements with no positional correlation are classified as not overlapping.

**Figure S5.**
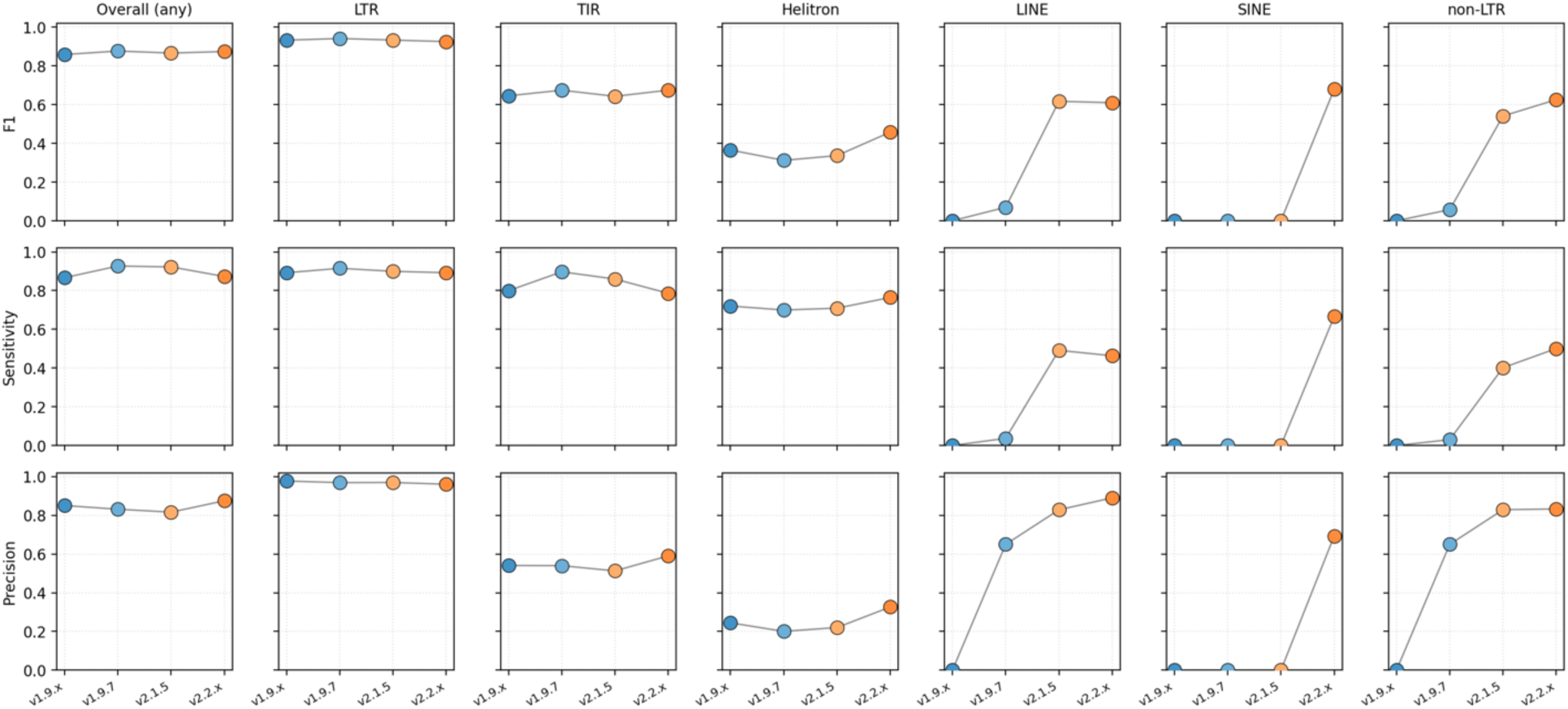
Per-category performance of EDTA releases on rice (MSU7). F1, sensitivity, and precision (rows) across overall, LTR, TIR, Helitron, LINE, SINE, and non-LTR categories (columns) for v1.9.x, v1.9.7, v2.1.5, and v2.2.x. No curated library is supplied in these runs. Performance on LTR, TIR, and Helitron is maintained across versions, while non-LTR detection (LINE, SINE, non-LTR) is substantially boosted in v2.

**Figure S6.**
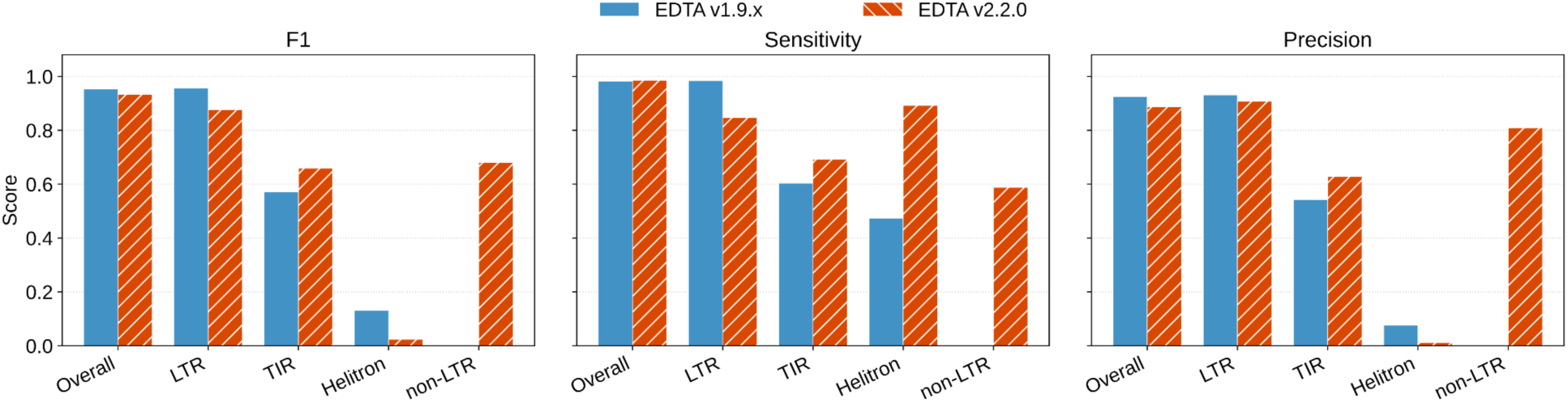
EDTA v2 maintains similar qualities of LTR, TIR, and Helitron annotations while extending non-LTR annotations on maize. No curated library is supplied in these runs.

**Figure S7.**
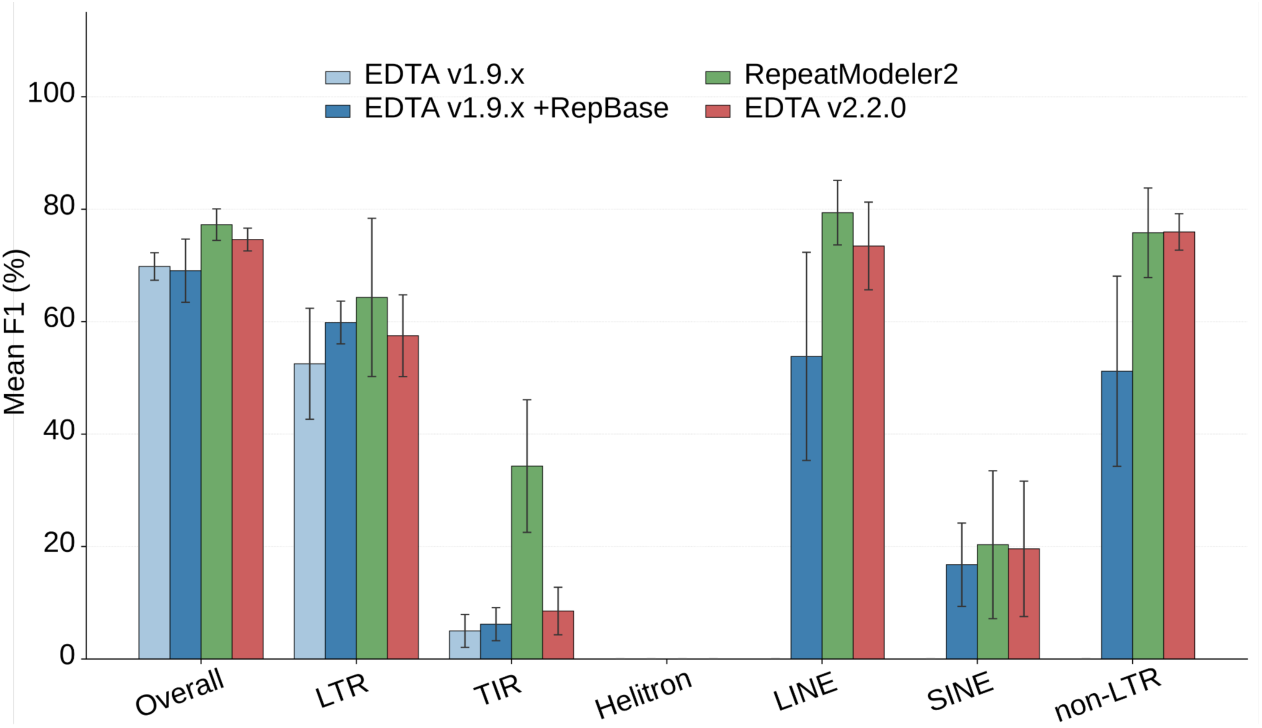
Mean F1 score on five animal models (chicken, fly, human, mouse, and zebra finch) for which curated TE libraries are available. All EDTA versions were run in *de novo*-only mode, except EDTA v1.9.x +RepBase, which was supplemented with curated TE libraries from RepBase. Error bars are standard deviation across species.

**Figure S8.**
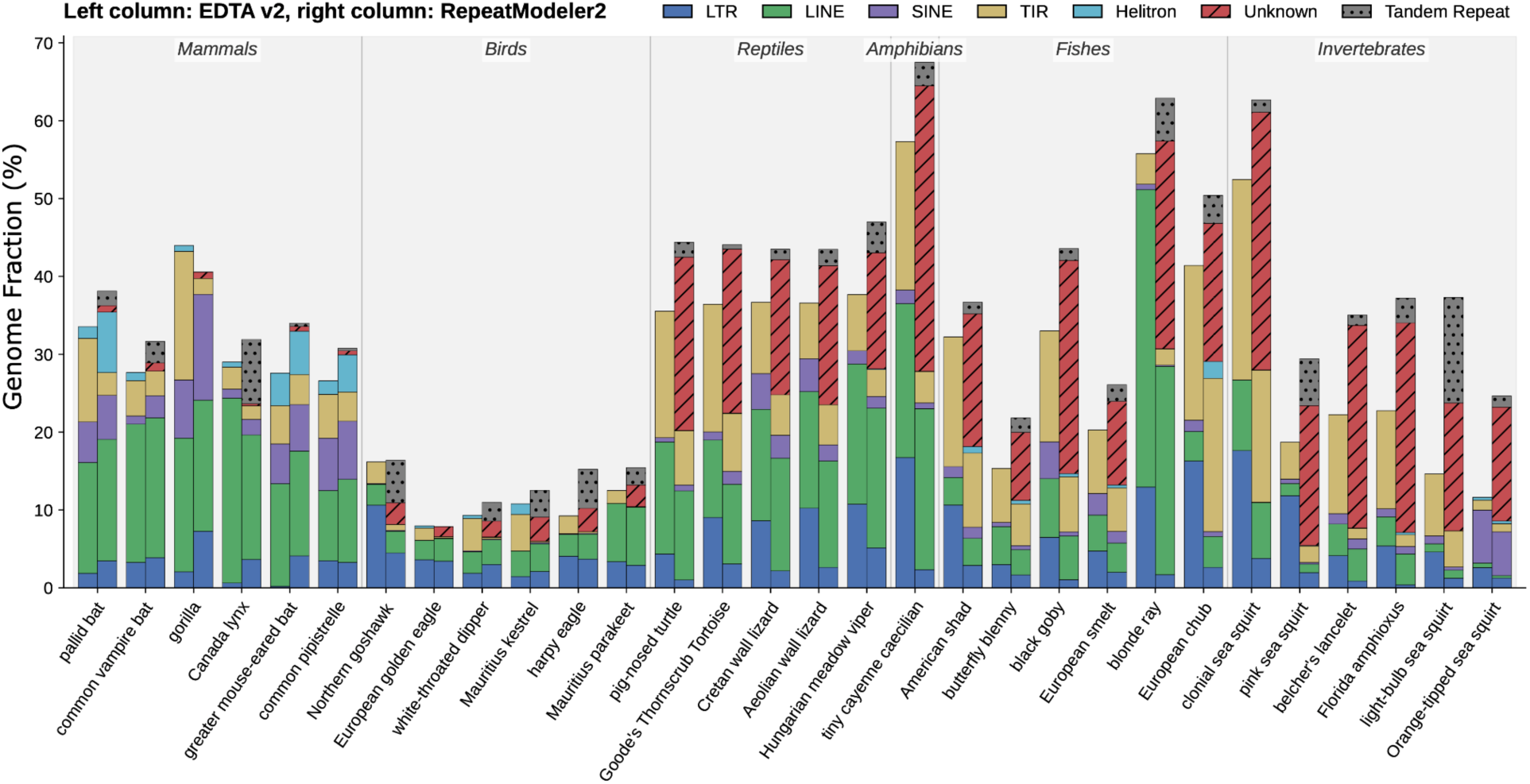
EDTA v2 resolves a larger fraction of repetitive DNA into named TE categories than RepeatModeler2 across 30 animal genomes. No curated library is supplied in EDTA v2 runs. For each species, paired stacked bars show the genome fraction (%) classified by EDTA v2 (left) and RepeatModeler2 (right) into five subclasses, the unclassified families (Unknown; red striped), and tandem repeats (grey dotted). Species are grouped along the x-axis by taxonomic class. Across most species, EDTA v2 annotates a comparable total TE fraction while assigning a substantially smaller fraction to the Unknown category.

**Figure S9.**
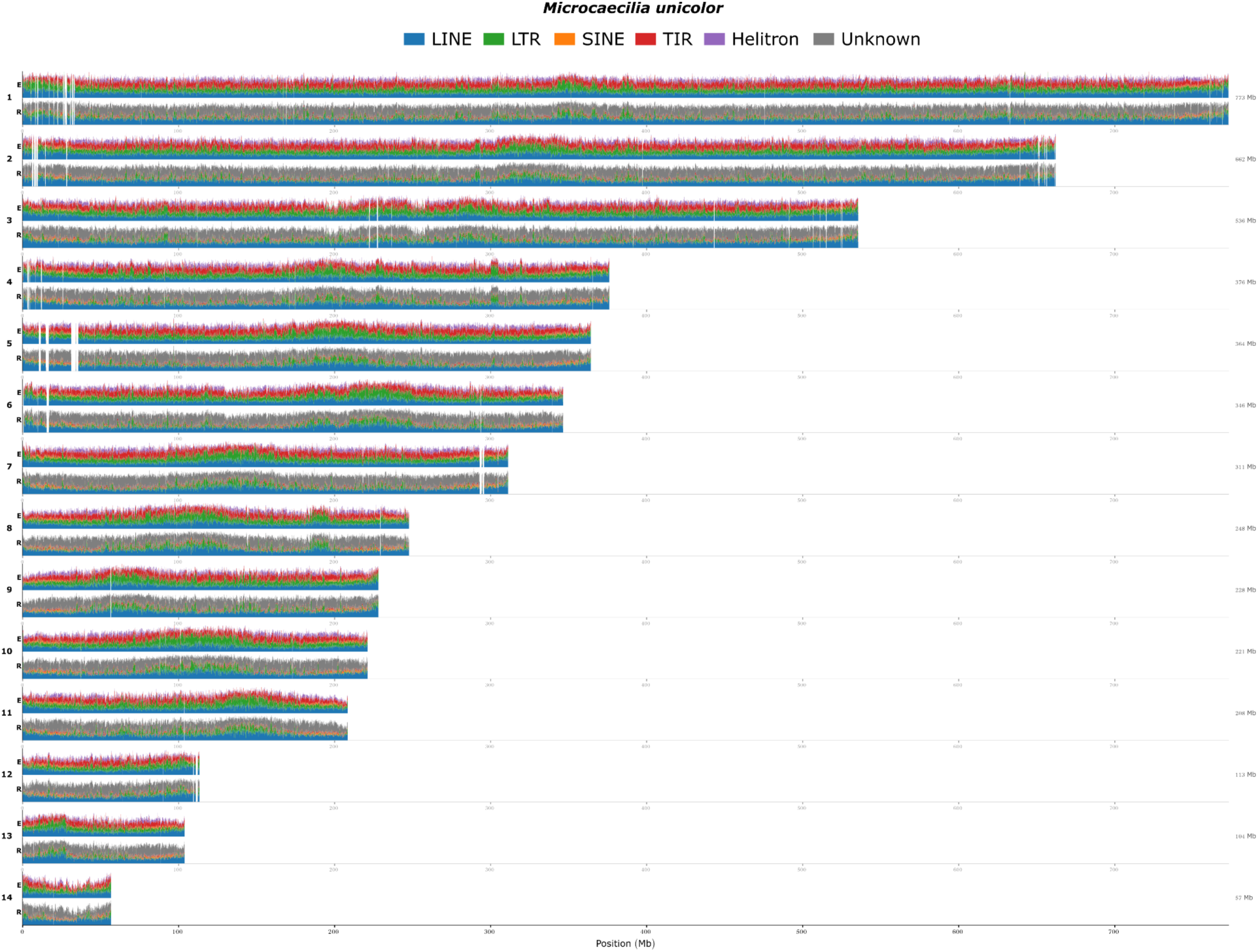
TE density across 14 chromosomes of *Microcaecilia unicolor*. EDTA v2 (E, top) and RepeatModeler2 (R, bottom) annotations are shown per chromosome in 100 kb windows. EDTA v2 annotates a larger fraction of classified TEs, with prominent contributions from TIR (red), LTR (green), and LINE (blue), whereas RepeatModeler2 carries a substantial Unknown component (gray).

**Figure S10.**
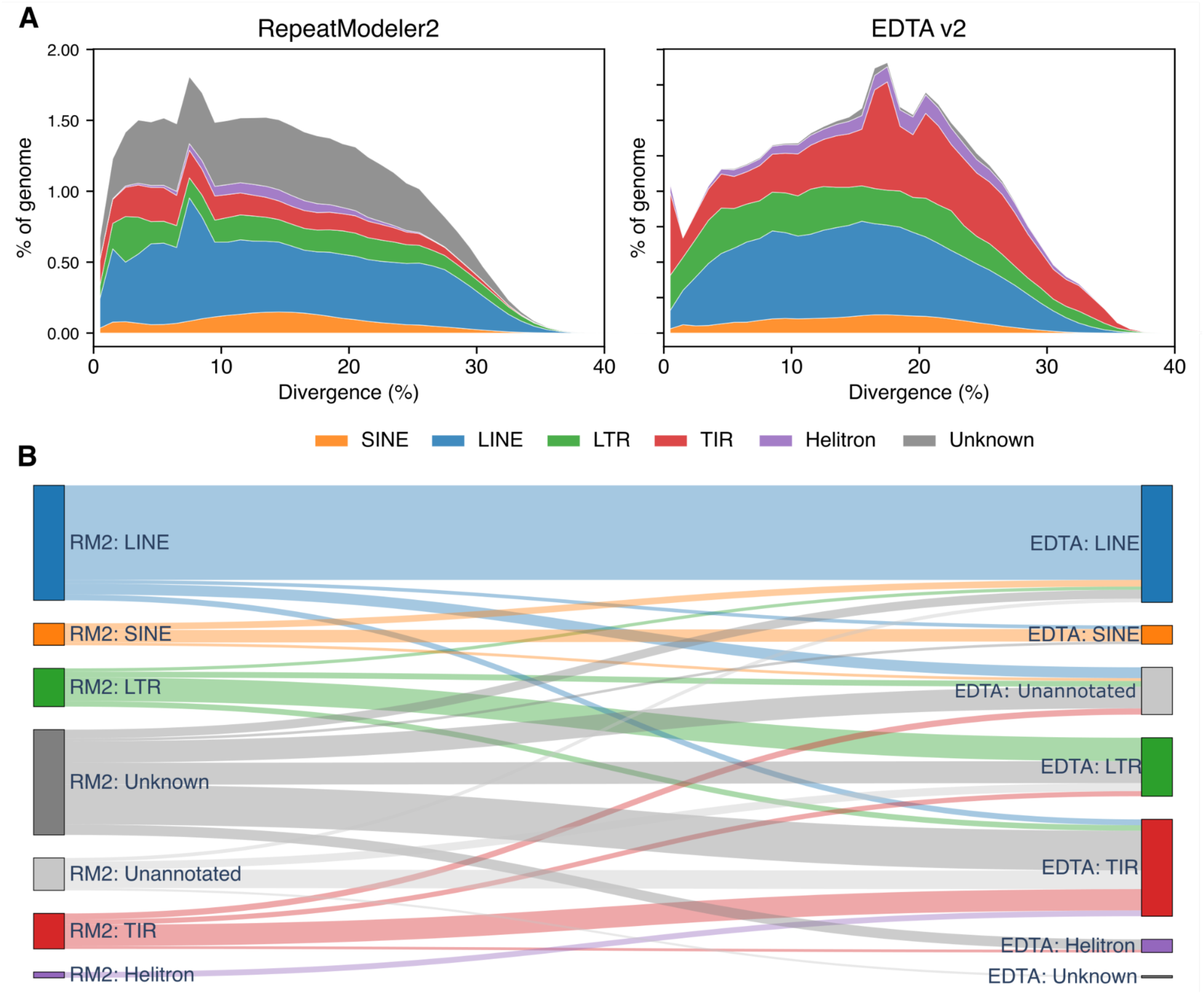
TE annotations between RepeatModeler2 and EDTA v2 across 30 VGP Phase 1 genomes. (**A**) Combined repeat divergence landscape across all 30 VGP species. Stacked areas show the proportion of genome (%) occupied by each TE subclass binned by substitution divergence from the library (1% bins). RepeatModeler2 leaves a substantial fraction of TEs unclassified (Unknown, gray), whereas EDTA v2 assigns most repeat content to specific subclasses, predominantly TIR and LTR elements. (**B**) Sankey plot of RepeatModeler2 (RM2) and EDTA v2 annotations across all 30 VGP species. Left nodes show RM2 classifications with tandem repeats purged from this summary; right nodes show EDTA v2 classifications for the same genomic intervals. Overlaps were at base pair resolution, with the flow width proportional to the number of base pairs (19.9 Gb total).

## Methods

### EDTA v2 pipeline overview

EDTA v2 processes a genome assembly through five sequential TE detection stages, performing *de novo* identification of LTR retrotransposons, SINEs, LINEs, TIR transposons, and Helitrons, each producing a raw library of candidate consensus sequences (**Fig. 1A**). These raw libraries pass through a cross-category purification processor (EDTA_processK) that resolves misclassifications between TE classes and reduces redundancy, then are concatenated into a non-redundant TE library. RepeatMasker v4.2.2 (ref^22^) uses this library to produce whole-genome TE annotations. Structurally intact TEs (LTR, TIR, and Helitron) are filtered and renamed by the library, producing whole-genome intact TE annotations. Compared with EDTA v1, which detected only LTR, TIR, and Helitron elements, v2 adds dedicated SINE and LINE detection stages, replaces the three-category filter with a five-category processor, and integrates LTR_HARVEST_parallel, HelitronScanner_parallel, and TIR-Learner v3 for accelerated structural TE identifications. EDTA v2 extends the checkpointing function of v1, which can automatically reuse existing intermediate results from time-consuming steps, so annotating large genomes on wall-time-limited facilities can be achieved.

### Genome sequence ID normalization

The namespace limitation (≤50 characters) of sequence IDs imposed by rmblastn inside RepeatModeler2 requires EDTA to limit sequence ID length to ≤15 characters, with 35 characters reserved for internal use, including adding sequence coordinates to the internal ID. In EDTA v1, ID shortening was arbitrary, which could create inconsistencies in downstream processing. EDTA v2 internally encodes sequence IDs using base62 characters within a 15-character space, then decodes internal IDs back to the originals at the end of the analysis. This scheme can encode 56.8 billion sequences in a single input file (maximum individual sequence length ≤1 Gb). For sequences between 1 Gb and 10 Gb, the maximum number of sequences encoded in a file is 238,328. For sequences longer than 10 Gb, users need to manually split them into segments < 10 Gb for EDTA v2 to process. This normalization is sufficient for large genomes (e.g., gymnosperms, amphibians), highly fragmented assemblies, and even FASTA files containing millions of raw sequencing reads.

### TE ID normalization

TE classification nomenclature varies widely across detection tools and reference libraries. For example, the hAT_TIR_transposon superfamily has been named as “hAT, hAT-Ac, Ac-Ds, Ac/Ds, DTA, DNA/DTA, DNAauto/hAT, DNAnona/hAT, DNA/hAT, TIR/hAT, DNA/hAT-Charlie, DNA/hAT-Tip100, DNA/hAT-Blackjack, DNA/hAT-hAT5, DNA/hAT-hAT6, DNA/hAT-hobo, DNA/hAT-Tol2, DNA/hAT-hATm-hAT-hybrid, …”, which creates inconsistencies and confusions in different methods and studies. To generate reproducible classification and annotation in the GFF3 output, EDTA v2 includes a curated Sequence Ontology (SO) lookup table that maps every classification alias to a canonical SO term and a stable SO accession. TE families lacking a dedicated SO entry are placed under the closest parent term. Regardless of which detector was originally used to name a TE, downstream files report the same SO term and ID for the same TE family. For the example above, all hAT aliases are renamed to hAT_TIR_transposon (SO:0002279).

### Accelerating LTR_HARVEST

LTR_HARVEST^23^ from GenomeTools v1.6.5 (ref^16^) is single-threaded and runs prohibitively long on large genomes. EDTA v2 wraps it in LTR_HARVEST_parallel v1.3, a multi-threaded Perl script that partitions the input genome into fixed-size pieces (default: 5 Mb) with a configurable overlap (default 100 kb). Each piece is dispatched to a worker thread running LTR_HARVEST independently. At the end of the run, per-piece candidate lists are concatenated, deduplicated against overlapping regions, and converted to whole-genome coordinates. To prevent simple-repeat-rich regions from stalling the pipeline, each thread is subject to a wall-clock timeout (default 120 s). On timeout, the offending piece is recursively re-split into 50-kb sub-regions and retried with LTR_HARVEST scans. The result is a faithful drop-in replacement that scales near linearly with the number of available cores.

### Accelerating HelitronScanner

HelitronScanner v1.1 (ref^24^) runs four sequential sub-commands (scanHead, scanTail, pairends, draw) in both strand orientations and is single-threaded. EDTA v2 wraps it in run_helitron_scanner.py, a Python multiprocessing script that partitions the genome into overlapping pieces (default: 10-Mb chunks with 50-kb overlap). The script then dispatches each chunk to a worker process that executes the full HelitronScanner workflow, and finally concatenates the outputs and converts them to whole-genome coordinates. Post-processing by format_helitronscanner_out.pl and filter_helitronscanner.pl produces output identical in format to the original serial run. The result scales near linearly with the number of available cores.

### Reimplementing TIR-Learner

TIR-Learner v2.5 (ref^25^) was a workflow orchestrating thirteen standalone Python scripts, passing intermediate state as files on disk. It required a brittle stack of Python 3.6, Keras 2.2.4, TensorFlow 1.14.0, scikit-learn 0.19.0, together with GenomeTools, GRF v1.0.2 (ref^17^), and utilities whose pinned versions conflicted across operating systems. Users routinely failed to install TIR-Learner on their production systems or existing conda environments. Furthermore, a large number of intermediate small files created heavy temporary I/O, prohibiting the run from staying within allocated system resources. Finally, the *de novo* TIR candidate discovery in TIR-Learner v2.5 relied solely on GRF, which scales poorly with fragmented assemblies.

We reimplemented TIR-Learner in v3, addressing these limitations through a full architectural rewrite and packaging overhaul. The workflow was replaced by a single TIRLearner Python class whose state is a dictionary of pandas DataFrames held in memory, with per-row operations parallelized via Swifter and a checkpoint system that writes timestamped intermediate information after every step for job recovery. The CNN model was migrated from TensorFlow 1 to Keras 3 with the PyTorch backend, significantly improving cross-platform compatibility and long-term maintainability. The new TIR-Learner is distributed via Bioconda so that all dependencies (Python ≥3.8, scikit-learn ≥1.3, Keras ≥3.3.3, PyTorch, GenomeTools, GRF, BLAST+) resolve through a single recipe.

TIR-Learner v3 adds TIRvish^16^ from GenomeTools (parameters: seed=20, mintirlen=10, maxtirlen=1000, similar=80, mintsd=2, maxtsd=11) as a second discovery engine. Candidate lists from TIRvish and GRF are concatenated and passed through the same filters. To accelerate GRF and TIRvish on fragmented assemblies, long sequences are split into overlapping 5-Mb windows (50 kb overlap) and processed in parallel. GRF’s native multithreading can be inefficient when inputs comprise many short sequences, where per-sequence loading overhead may exceed the actual analysis time. We therefore implemented a hybrid parallelization scheme that combines native threading with an outer multiprocessing layer to mitigate this (**Fig. S2, S3**). Both overlapping windows and the use of multi-search engines (GRF and TIRvish) yield duplicate candidates; therefore, results are deduplicated by sorting all candidates by genomic coordinate and length, then comparing adjacent entries for positional overlap between their main bodies and TIRs. Among partially overlapping candidates, the shorter element is discarded, whereas fully nested or non-overlapping candidates are retained as distinct elements (**Fig. S4**). EDTA v2 invokes the TIR-Learner v3 pipeline directly, and the resulting raw library is handed to TEsorter v1.5.x cleanup before entering the cross-category filter.

### Non-LTR detection

In EDTA v1, non-LTR detection was available only via the “--sensitive 1” parameter, which invoked RepeatModeler2 at a late filtering stage. At this stage, non-LTR candidates were identified from the remaining sequences after masking with the filtered TE library. This strategy could fail if the filtered library contained non-LTR contaminants that eliminate non-LTR candidates in the masking step. In EDTA v2, non-LTR detection is promoted to the same level as other TE subclasses. RepeatModeler2 is run *de novo* on the genome using the NCBI engine, and its output is processed by TEsorter for purging class-mismatched sequences. Sequences assigned as LINE or SINE are retained; the non-LINE/SINE residue is retained as a supplementary library used when “--sensitive 1” is specified. To further enhance SINE annotation, AnnoSINE_v2 (ref^14^) is run with parameters “-a 2 – num_alignments 50000 –copy_number 3 –shift 100 –auto 1 3”. A family is retained only if at least three genomic copies are recovered.

### Per-library classification cleanup with TEsorter

Before cross-category filtering, each of the five raw libraries is independently cleaned of misclassification by TEsorter v1.5.x (ref^15^), which classifies sequences using profile hidden Markov models for conserved TE protein domains. The “--disable-pass2” parameter retains only the strongest pHMM evidence without homology classification of unclassified TEs. The cleanup_misclas.pl script rejects sequences whose original subclass and superfamily disagree with TEsorter’s classification.

### Cross-category library filtering using richness ratio

Raw libraries from all five detection stages are filtered through EDTA_processK.pl, replacing the three-category processor in EDTA v1. Filtering proceeds in three phases. First, TE candidates were compared across groups using the richness ratio to remove cross-contamination between subclasses. For example, LTR, TIR, and Helitron candidates were compared against one another, and regions more enriched in another TE type were masked and removed. Richness ratio cutoffs were set to 1 for LTR and TIR candidates and 1.5 for Helitrons, because Helitron predictions often include neighboring non-Helitron sequences. Second, RepeatMasker was used for additional homology-based cleaning (divergence ≤ 40%) following the hierarchical order. To begin, LINE sequences were used to clean LTR candidates. Then, cleaned LTR plus LINE sequences were used to clean SINE candidates. Finally, LTR, SINE, and LINE sequences were used to clean TIR and Helitron candidates. Third, remaining redundancy was removed from candidates using all-vs-all BLAST clustering with cleanup_nested.pl, requiring at least 95% query coverage and prioritizing the removal of shorter redundant sequences.

### Purging nested TEs

Nested TE removal is performed by cleanup_nested.pl with all-versus-all BLASTs of the input library and removes candidates detected as substrings of other candidates. The original implementation in EDTA v1 considered only the first BLAST high-scoring pair (HSP) per search. When homologous elements were separated into several short, non-overlapping HSPs, they were not recognized as nested copies and persisted as redundant entries. EDTA v2 collects every HSP between a query and subject, merges overlapping HSPs, and sums total non-overlapping coverage before applying the –cov cutoff. In test libraries, this change reduced redundant entries by 10-20% without measurable impact on annotation sensitivity or specificity.

### Reconstructing fragmented TEs

RepeatMasker can produce fragmented annotations due to high penalties on gap opening in the BLAST search step. The combine_RMrows.pl script iteratively merges fragmented hits deriving from the same physical TE based on these criteria: 1) adjacent entries are merged when they share chromosome, strand, element name, class, subclass, and superfamily; 2) the genome gap is ≤35 bp; 3) divergence values differ by ≤3.5%; and 4) TE-coordinate offsets are continuous (≤35-bp gap on the library sequence axis). Merged rows use length-weighted averages of SW_score, div, del, and ins in the result.

### Plotting TE landscapes

EDTA v2 emits summary figures alongside the GFF3 when –anno 1 is set. Two visualizations are generated: (1) a divergence plot summarizing Kimura two-parameter distances between annotated copies and library sequences, displaying relative ages of each TE class as histograms; and (2) a chromosomal density plot computing per-superfamily TE coverage in 1-Mb sliding windows (500-kb step), rendering class-specific banding patterns along each chromosome.

### Benchmarking accelerated detectors

TIR-Learner versions were benchmarked across a range of genome sizes. Implementations tested: (i) TIR-Learner 2.5, (ii) TIR-Learner 3.0 with TIRvish disabled (-a skip_tirvish), and (iii) TIR-Learner 3.0 with TIRvish enabled (default). The test genomes were three subsets of the highly fragmented Pacific white shrimp (*Penaeus vannamei*) genome^21^ at 312 Mb (contig N50 70.5 kb), 640 Mb (contig N50 80.9 kb), and 1.62 Gb (contig N50 87.4 kb). All runs used identical parameters (-l 5000 –s other, 28 threads) on the same compute node.

HelitronScanner scalability was evaluated by benchmarking the parallelized wrapper (run_helitron_scanner.py) against the original single-threaded wrapper (run_helitron_scanner.sh) on seven genomes spanning 312 Mb to 29.1 Gb: axolotl (*Ambystoma mexicanum*, GCF_040938575.1, 29.1 Gb), fire-bellied toad (*Bombina bombina*, GCF_027579735.1, 10.0 Gb), horn shark (*Heterodontus francisci*, GCF_036365525.1, 6.0 Gb), clouded leopard (*Neofelis nebulosa*, GCF_028018385.1, 2.5 Gb), and *P. vannamei* (GCA_003789085.1; 1.6-Gb full assembly plus 640-Mb and 312-Mb subsets). The parallel implementation benchmark used 128 CPUs on a single node.

### Benchmarking EDTA v2 annotation quality

EDTA v2 performance was benchmarked on seven reference genomes: rice (*Oryza sativa* MSU7), maize (*Zea mays* B73 v5), *Drosophila melanogaster* (r6.28), chicken (*Gallus gallus*, galGal6), zebra finch (*Taeniopygia guttata*, taeGut2), mouse (*Mus musculus*, mm39), and human (*Homo sapiens*, hg38). For the four vertebrate genomes, ground-truth annotations were UCSC RepeatMasker tracks obtained from hgdownload.soe.ucsc.edu/goldenPath. For the remaining genomes, reference annotations were generated via RepeatMasker with curated species-specific libraries: a manually curated *D. melanogaster* library (https://github.com/bergmanlab/drosophila-transposons), the non-redundant rice TE library shipped with EDTA, and the Maize TE Consortium library (https://github.com/oushujun/MTEC)^26^. A limitation is noted: the UCSC vertebrate tracks reflect RepBase snapshots from 2013-2018 and may under-call subsequently curated TE families.

Annotation quality was measured with lib-test.py, a Python rewrite of lib-test.pl, a script developed in the original EDTA. Both implementations compute sensitivity, specificity, accuracy, precision, FDR, and F1 across eight TE categories (LTR, nonLTR, LINE, SINE, TIR, MITE, Helitron, and total). The rewrite replaces BED-based operations with in-memory bitarray operations, producing all categories in one pass with much accelerated performance (evaluation generally finished in 1-2 minutes compared to 30 minutes in the original version). The Python version additionally generates a superfamily-by-superfamily confusion matrix for quality assessments.

### Annotation inconsistency evaluation

We evaluated the annotation inconsistency of each whole-genome TE annotation using evaluation.pl from the EDTA package. For a given RepeatMasker.out annotation file, the pipeline (1) extracts all annotated TE sequences with Smith-Waterman score > 300 and alignment length > 80 bp from the genome assembly; (2) performs an all-versus-all BLASTn search among extracted sequences (word size 7, e-value 1e-5, minimum identity 80%, minimum alignment length 80 bp); and (3) for each pair of homologous sequences, records whether the two carry the same or different subclass labels. Then, a confusion matrix is built across TE subclasses, with the per-subclass inconsistency rates computed as the fraction of pairwise hits in which the partner carries a different subclass label. A higher inconsistency rate indicates that homologous TE copies in the genome were assigned conflicting subclass labels, reflecting annotation unreliability without relying on curated reference annotations.

### TE annotation across 30 animal species

A representative set of 30 animal species was collected: 6 mammals, 6 birds, 6 fish, 5 non-avian reptiles, 1 amphibian, and 6 invertebrates (**Table S1**). A phylogenetic tree of these species was obtained from the Open Tree of Life (OToL)^27^. The rectangular cladogram with TE composition bars was plotted using matplotlib^28^. EDTA v2.2.0 annotated all species with default parameters and without curated libraries. TE content was quantified at the subclass level (LTR, LINE, SINE, TIR, Helitron, and Unknown) as a fraction of total assembly size.

RepeatModeler2 annotations (v2.0.4, default parameters) were obtained from the UCSC genome browser. Species-specific tandem repeat (TR) libraries were generated using the satellome v1.3.0 pipeline (https://github.com/aglabx/satellome), which runs Tandem Repeat Finder (TRF) v4.09 (ref^29^) across each genome assembly and applies a series of copy-number filters to distinguish *bona fide* satellite sequences. Candidate arrays were retained if their monomer period was at least 10 bp (excluding microsatellites), the tandem array spanned at least 100 bp, and the within-array copy number was at least 50, ensuring that retained sequences represent abundant, genuinely repetitive satellites rather than isolated tandem duplications. Monomer consensuses were then clustered at 80% identity using cd-hit-est. Finally, TEsorter was used to remove any monomers carrying recognizable TE protein domains. The resulting TR libraries were used to mask each species’ RepeatModeler2 consensus sequences with RepeatMasker v4.2.2, identifying which positions in each RM2 consensus overlapped TR content. These TR coordinates were then cross-referenced with the genome-wide RepeatModeler2 annotations to proportionally split the original annotation into TE and TR components. Across the 30 species, TR content ranged from 0.01% (*Aquila chrysaetos*) to 13.56% (*Clavelina lepadiformis*).

## Notes

### Competing Interest Statement

The authors have declared no competing interest.

